# Optimization of the *Ds*Red fluorescent protein for use in *Mycobacterium tuberculosis*

**DOI:** 10.1101/383836

**Authors:** Paul Carroll, Julian Muwanguzi-Karugaba, Tanya Parish

## Abstract

**ABSTRACT:** *Objective:* We have previously codon-optimized a number of red fluorescent proteins for use in *Mycobacterium tuberculosis* (mCherry, tdTomato, Turbo-635). We aimed to expand this repertoire to include DsRed, another widely used and flexible red fluorescent protein.

*Results:* We generated expression constructs with a full length *DsRed* under the control of one of three strong, constitutive promoters (P_hsp60_, P_rpsA_ or P_G13_) for use in mycobacteria. We confirmed that full length DsRed (225 amino acids) was expressed and fluoresced brightly. In contrast to mCherry, truncated versions of DsRed lacking several amino acids at the N-terminus were not functional. Thus, we have expanded the repertoire of optimized fluorescent proteins for mycobacteria.

## INTRODUCTION

Fluorescent proteins (FPs) have become the work horses of molecular biology and microbiology, with numerous applications. A plethora of variants of *Aequorea victoria* green fluorescent protein (GFP) (1) and *Discosoma* sp red fluorescent protein (*Ds*Red) (2) are available covering almost the whole light spectrum from green to infra-red (3). Mutant derivatives have been engineered with altered excitation and emission wavelengths, increased or decreased stability, resistance to photo bleaching, sensitivity to environmental stimuli and substrates, as well as time for fluorophore maturation, intrinsic brightness and multimeric formats (3, 4). We previously described the use of a range of red reporters, of which the brightest was mCherry (5). We wanted to expand our repertoire of proteins. Since DsRed has been widely used as a bright and stable reporter, we optimized constructs for its expression in *M. tuberculosis.*

## MAIN TEXT

### MATERIALS AND METHODS

#### Bacterial culture

*Escherichia coli* DH5α was cultured in LB medium or on LA agar. *M. tuberculosis* H37Rv was grown in Middlebrook 7H9 medium plus 10% v/v OADC (oleic acid, albumen, dextrose, catalase) supplement (Becton Dickinson) and 0.05% w/v Tween 80 or on Middlebrook 7H10 agar (Becton Dickinson) plus 10% v/v OADC. Hygromycin was used at 100 μg/ml where required.

#### Construction of expression vectors

The *Ds*Red expression vectors were constructed as follows: a partial *Ds*Red sequence was codon optimized for *M. tuberculosis,* synthesised and cloned into pUC57 (Genscript USA Inc.) to generate pRed1. The *Ds*Red ORF was excised from pUC57 as a BamHI/HindIII fragment and cloned into pSMT3 (6) to generate pBlaze1. The *Ds*Red ORF was extended three times by PCR to generate pRedA1, pRedB1 and pRedC1 using primers DsRed-F1 5’- <u>GGA TCC</u> **ATG** CGC TTC AAG GTG CGC ATG GAG GGC TCG GTG AAC-3’, DsRed-F2 5’- <u>GGA TCC</u> **GAC** GTG ATC AAG GAG TTC ATG CGC TTC AAG GTG CGC-3’ and DsRed-F3 5’- <u>GGA TCC</u> **ATG** GCC TCG TCG GAG GAC GTG ATC AAG GAG TTC together with the reverse primer *Ds*Red-R 5’- <u>AAG CTT</u> TTA CAG GAA CAG GTG GTG CCG-3’. The restriction sites are underlined, potential start codons are in bold. The ORFs were excised and cloned into pSMT3 (6) as *Bam*HI/ *Hin*dlll fragments to generate pBlazeA1, pBlazeB1 and pBlazeC1 with *Ds*Red under the control of the *hsp60* promoter. Plasmids pBlazeC8 and pBlazeC10 were generated by replacing P_hsp60_ with P_rpsA_ and P_G13_ respectively. All three promoters should drive constitutive high level expression (5, 7, 8).

#### Quantitation of fluorescence in whole cells

*M. tuberculosis* was electroporated as described (9) and transformants selected with hygromycin. *M. tuberculosis* was grown to stationary phase, harvested, washed twice in 10 mM Tris pH 8.0 and resuspended in 10 mM Tris pH 8.0 to an OD_580_ of 0.25, 0.10, 0. 05 and 0.01 in 12 × 100 mm glass culture tubes. Fluorescence was measured on a Shimadzu RF-1501 spectrofluorimeter (Shimadzu) with a detection range of 0-1015 relative fluorescent units at Ex/Em 558/583nm (5).

#### Western analysis of fluorescent proteins

Cell extracts were prepared from liquid cultures. Cells were harvested by centrifugation, washed twice in 10□mM Tris (pH 8.0), resuspended in 1□ml of 10□mM Tris (pH 8.0), and added to lysing matrix B tubes (QBiogene). Cells were disrupted using the Fastprep (QBiogene) set at speed 6.0 for 30 seconds. Samples were centrifuged for two min, and the supernatant was recovered and filter sterilized. Protein was quantified using a BCA kit (Pierce), and 10□μg of total protein was subjected to Western blot using a rabbit anti-body (Clonetech). The primary antibody was detected using horseradish peroxidase goat-anti-rabbit (Sigma), and activity was detected using an ECL kit (GE Healthcare).

## RESULTS

We were interested in the use of FPs in *M. tuberculosis* and had previously used these as reporters of bacterial viability for *in vitro* and *in vivo* studies (5, 8). We were successful in obtaining high level expression by using codon-optimized versions of red fluorescent proteins driven by strong mycobacterial promoters (5).

### Optimization of *Ds*Red expression

We wanted to expand the range of reporters available for use to increase flexibility and allow dual reporter expression and monitoring. We selected *Ds*Red for optimization, based on its Ex/Em wavelengths, and the fact that it is a well-characterized FP in wide use.

### Expression of *Ds*Red uses a different translational start site than mCherry

Our initial attempts to obtain expression of a codon-optimized *Ds*Red were unsuccessful. We constructed a synthetic gene for *Ds*Red using a similar approach as we used with another red fluorescent protein mCherry (Figure 1). We designed a codon-optimized version based on the *Ds*Red-T3 protein previously used. We cloned the synthetic version into a mycobacterial expression vector and tested for fluorescence in *M. tuberculosis*. Surprisingly, we did not detect any fluorescence from this construct (Figure 1C).

**Figure 1.**
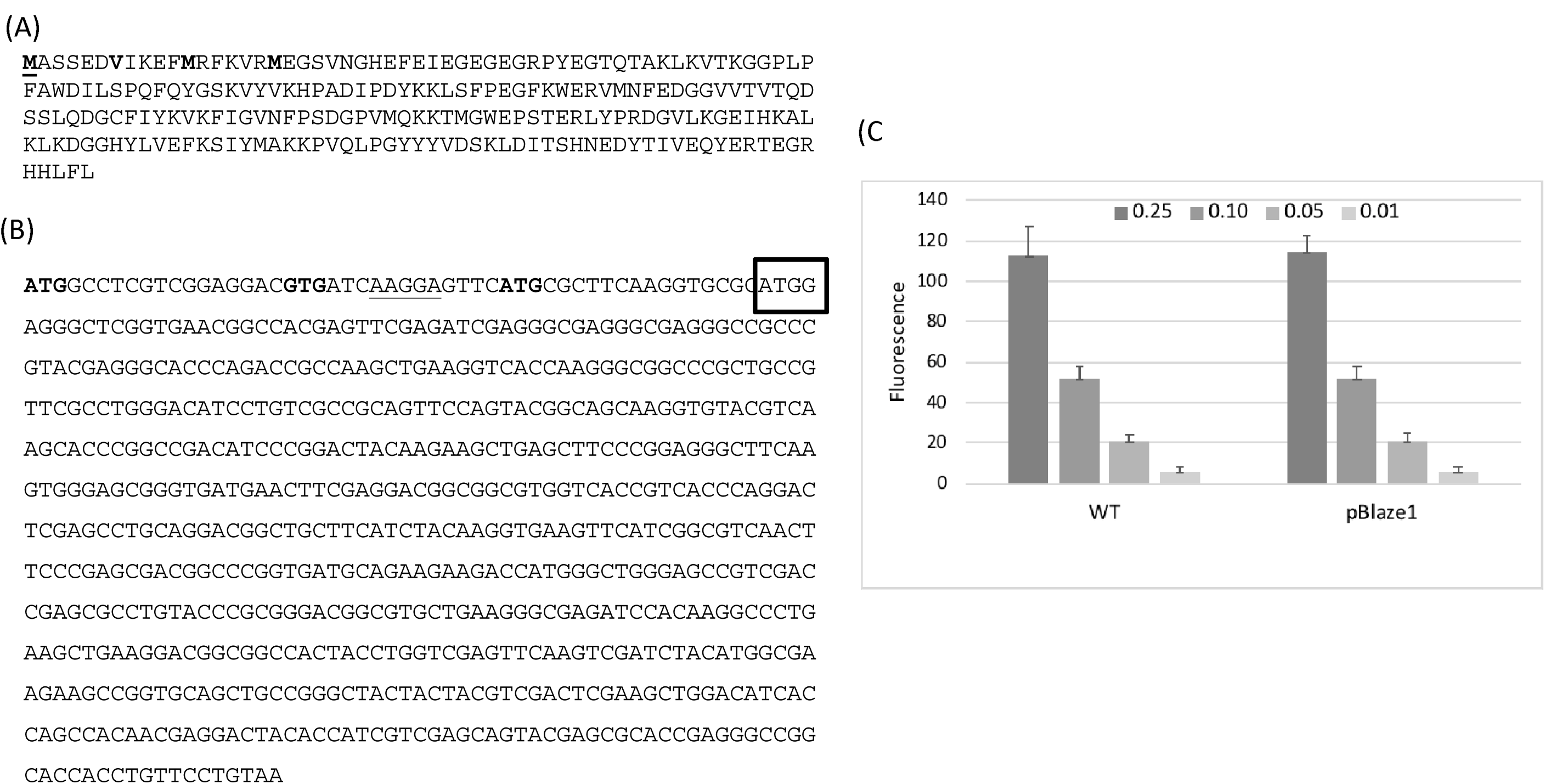
Expression of non-functional *Ds*Red. **(A)** *Ds*Red Protein sequence. Three potential translational start sites (methioinine) are indicated in bold. The valine which corresponds to the methionine start site of mCherry in *M. tuberculosis* is also indicated in bold. **(B)** DNA sequence of *Ds*Red. The 5’ end of the synthetic gene designed to codon-optimize *Ds*Red for *M. tuberculosis* is boxed. Potential starts sites are indicated in bold. The Shine Delgarno sequence is underlined. (C) *M. tuberculosis* was resuspended in 10 mM Tris pH 8.0 to an OD_580_ of 0.25, 0.10, 0.05 and 0.01 in 12 × 100 mm glass culture tubes. Fluorescence was measured at Ex/Em 558/583nm. WT - wild-type (no plasmid). pBlaze1 – recombinant strain carrying *Ds*Red 208aa. Data are the average ± SD of three cultures.

mCherry is a variant of *Ds*Red and we expected the two proteins would be similarly functional. Our previous work demonstrated that mCherry is expressed from a distal translational start site than the one annotated in the databases (10). Sequence alignment shows the few mutations which differ between the two (Figure 2A); these include loss of the translational start site we identified for mCherry, although there are still multiple translational start sites (Figure 1A). The version we used for the synthetic gene used a downstream translation start site and would produce a truncated version of *Ds*Red as compared to mCherry. Therefore it was possible that we did not express the full protein (Figure 1B). In order to determine the functional start site for *Ds*Red we used a different approach in which we cloned several versions of the coding region into the expression vector under the control of the constitutive *hsp60* promoter (Figure 2B).

**Figure 2.**
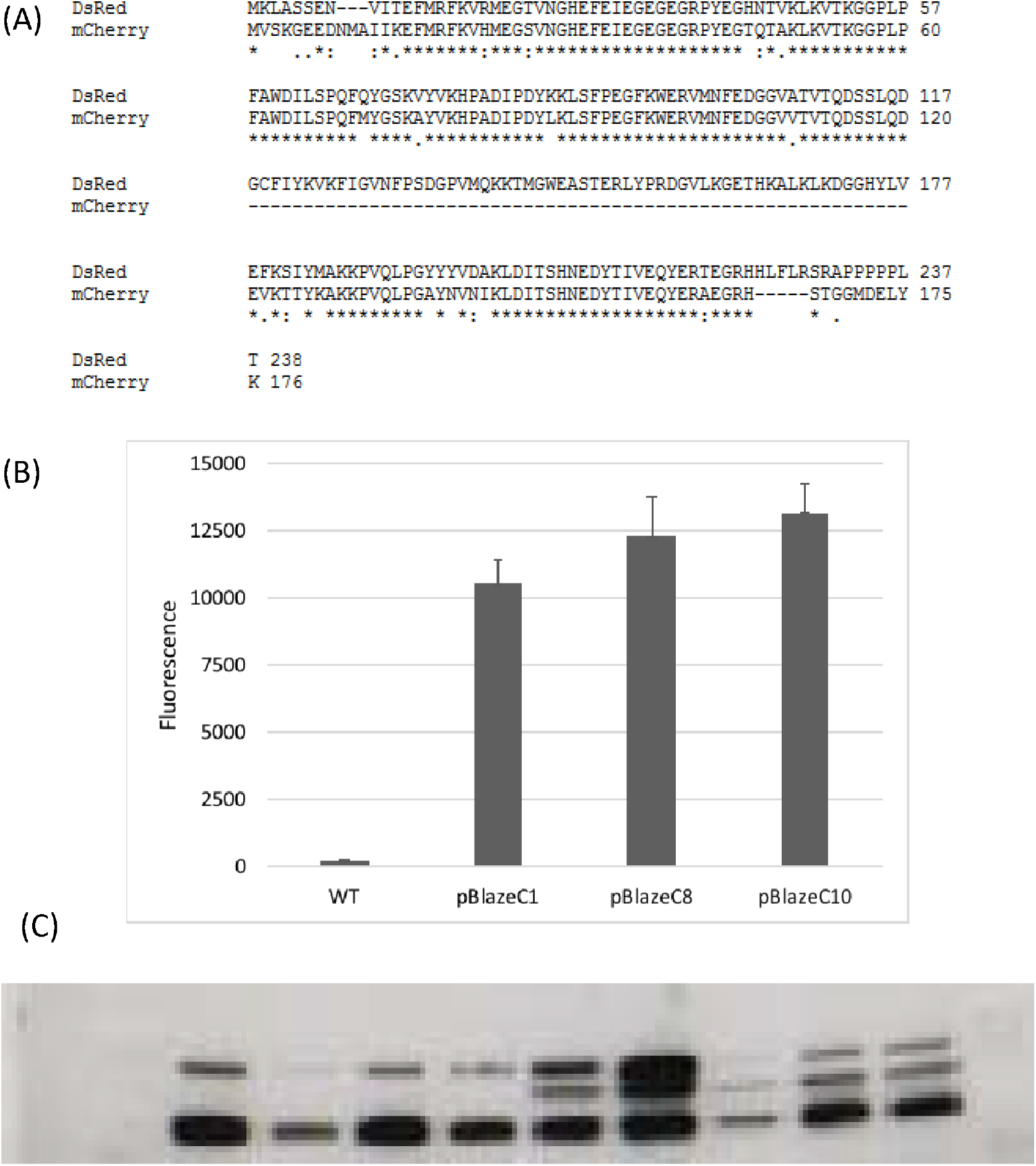
Expression of functional *Ds*Red. **(A)** Sequence alignment of mCherry and *Ds*Red. Protein sequences were aligned using Clustal (13). **(B)** Activity of full length *Ds*Red expressed from mycobacterial promoters. pBlazeC1 – P_hSp60_; pBlazeC8 - P_rpsA_; pBlazeC10 – P_G13_. Recombinant *M.tuberculosis* was resuspended in 10 mM Tris pH 8.0 to an OD_580_ of 0.25, 0.10, 0.05 and 0.01 in 12 × 100 mm glass culture tubes. Fluorescence was measured at Ex/Em 558/583nm. Data are the average ± SD of three cultures. **(C)** Plasmids were transformed into *E. coli* and cell-free extracts analyzed by Western blotting; 10 μg protein were subjected to SDS-PAGE, blotted onto PVDF membrane and visualized with anti-*Ds*Red antibody. Lane 1s and 11 – empty; Lane 2 – *E. coli* (no plasmid); Lane 3 – pRed1; Lane 4 – pRedA1; Lane 5 – pRedB1; Lane 6 – pRedC1; Lanes 7 and 8 – pBlazeC1; Lane 9 - pBlazeC8; Lane 10 – pBlazeC10. The arrow indicates the size of the *Ds*Red protein.

**Table 1.**
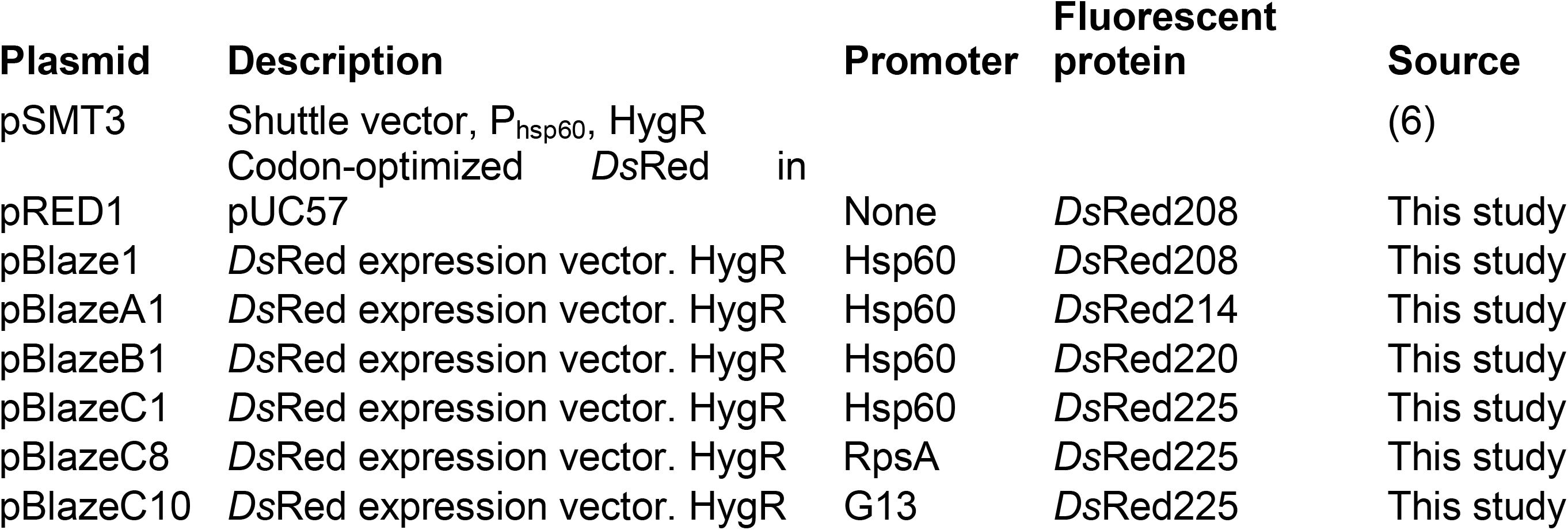
Plasmids used in this study.

In order to test this, we used PCR amplification to extend the region sequentially. We extended the gene to incorporate both additional start sites and generate proteins of 214, 220 and 225 amino acids. These variants were cloned into the same mycobacterial expression system and tested. Plasmids were transformed into *M. tuberculosis* and fluorescence was monitored. In contrast to mCherry, expression of a functional fluorescent *Ds*Red was not seen with any truncated versions of the protein. In fact fluorescence could only be detected when the full length amino acid sequence (as annotated) was cloned into the expression vector; high level fluorescence was seen with transformants carrying the plasmid pBlazeC1 (Figure 2B).

We constructed two alternative vectors with *Ds*Red under the control of either P_rpsA_ or P_G13_ (pBlazeC8 and pBlazeC10 respectively); both of these constructs gave high level expression in *M. tuberculosis*. Western blotting using an anti-*Ds*Red antibody in *E. coli* demonstrated that a protein of the expected size was only seen in bacteria carrying the full length construct (pBlazeC series), but not in the strains carrying the truncated version (Figure 2C).

## DISCUSSION

We have determined that the functional translational start sites for two closely related FPs are different in *M. tuberculosis*. Although mCherry was functional even when a truncated version was being expressed, *Ds*Red was non-functional in a truncated form and only fluoresced when expressed as a full length protein (225 amino acids). Western blotting suggested that the lack of fluorescence was most likely due to a lack of protein expression, since proteins could not be detected in the plasmids carrying truncated forms. This difference may relate to protein stability, with the extended N-terminal portion of *Ds*Red increasing stability or protein maturation; alternatively this could be attributed to the physical state of the active proteins, since mCherry functions as a monomer, whereas *Ds*Red is a tetramer which might also affect protein degradation.

In conclusion, we have codon-optimized *Ds*Red for use in *M. tuberculosis* and demonstrated its high level fluorescence in that species from three different promoters of slightly varying strength (*hsp60, rpsA,* and *G13*). These vectors extend our current repertoire of functional fluorescent proteins for mycobacteria. They will be useful for generating fluorescent strains of *M. tuberculosis* for use in multiple studies, such as monitoring drug efficacy *in vitro* and *in vivo* (5, 8, 11, 12) and will allow for detection of multiple reporters simultaneously.

### Limitations

- We have monitored the expression of *Ds*Red under aerobic conditions only.
- We have not monitored long term stability of expression in the absence of antibiotic selection to maintain the plasmid.
- We have not monitored stability of expression *in vivo*.

## Declarations

### Ethics approval and consent to participate

Not applicable

### Availability of data and materials

All data generated or analysed during this study are included in this published article.

## Acknowledgements

We thank Amanda Brown and Lise Schreuder for technical assistance and helpful discussion.

## Consent for publication

Not applicable

## Author’s contributions

Experimental design: PC, JM, TP

Experimental work: PC, JM

Data analysis: PC, TP

Writing manuscript: PC, TP

Reviewing manuscript: PC, JM, TP

All authors have read and approved the manuscript.

## Abbreviations

FP: fluorescent protein
OADC: oleic acid, albumin, D-glucose, catalase.

